# Yield-Stress and Creep Control Depot Formation and Persistence of Injectable Hydrogels Following Subcutaneous Administration

**DOI:** 10.1101/2022.04.20.488959

**Authors:** Carolyn K. Jons, Abigail K. Grosskopf, Julie Baillet, Jerry Yan, John H. Klich, Eric A. Appel

**Affiliations:** Department of Materials Science & Engineering, Stanford University, Stanford, CA 94305, USA; Department of Chemical Engineering, Stanford University, Stanford, CA 94305, USA; Department of Bioengineering, Stanford University, Stanford, CA 94305, USA; Stanford ChEM-H Institute, Stanford University, Stanford, CA 94305, USA; Woods Institute for the Environment, Stanford University, Stanford, CA 94305, USA; Department of Pediatrics - Endocrinology, Stanford University School of Medicine, Stanford, CA 94305, USA

**Keywords:** Hydrogels, Drug Delivery, Rheology, Therapeutics, Polymers

## Abstract

Hydrogels that can be injected into the body using standard needles or catheters enable a minimally invasive strategy to prolong local delivery of therapeutic drug and cellular cargo. In particular, physically crosslinked hydrogels exhibit shear-thinning and self-healing behaviors enabling facile injectability and depot formation upon administration. While prior efforts to characterize these systems have focused on injectability and cargo release behaviors, prediction of cargo release in the body often assumes the materials form a depot rather than spreading out upon administration. Here, we evaluate how hydrogel rheology correlates with depot formation and persistence following subcutaneous administration in mice with two physicochemically-distinct, physically crosslinked hydrogel systems. We evaluate calcium-alginate and polymer-nanoparticle hydrogel systems exhibiting variable mechanical behaviors across several rheological properties (stiffness, viscoelasticity, yield stress, and creep). By relating measured rheological properties to depot formation and persistence time following subcutaneous administration, we identify that yield stress is predictive of initial depot formation while creep is predictive of depot persistence. Indeed, only materials with yield stresses greater than 25 Pa form robust depots and reduced creep correlates with longer depot persistence. These findings provide predictive insights into design considerations for hydrogel technologies capable of extended controlled release of therapeutic cargo.

## 1. Introduction

Drug delivery is a pressing problem with the global controlled pharmaceutical delivery market projected to exceed USD 2.2 trillion by 2026.^[1]^ Hydrogels are a particularly exciting controlled delivery platform that are characterized by cross-linked macromolecular networks retaining a significant amount of water and which have been utilized for a range of translational applications including immunology,^[2,3]^ oncology,^[4,5,6]^ cardiology,^[7,8,9]^ tissue engineering,^[10,11]^ wound healing,^[12,13,14,15]^ and pain management.^[16]^ Their high water content is advantageous as it provides physicochemical similarity to biological tissues and allows for the encapsulation of therapeutic cells and hydrophilic drug cargo, while the presence of a polymer network imparts physical structure, tunable mechanical properties, and controlled cargo diffusion or cell motility. ^[17,18,19]^

Hydrogels allow for control over how drugs are available to cells and tissues over time and in space. Greater control over spatial and temporal drug delivery allows for improved therapeutic outcomes by enhancing treatment efficacy while reducing toxicity and required dosage.^[20,21,22]^ Hydrogels that can be injected into the body using standard needles or catheters enable a minimally invasive strategy to prolong local delivery of therapeutic cargo, whether cells or pharmaceuticals. In particular, physically-crosslinked hydrogels exhibit shear-thinning and self-healing behaviors enabling facile injectability and self-healing upon administration.^[ 23, 24, 25, 26 ]^ Subcutaneous administration is often a preferred means of drug administration as it is sufficiently simple to even allow for facile self-administration by patients. The modeling of release kinetics from a polymeric drug delivery system in the body is often simplified by assuming the formation of a spherical depot following subcutaneous administration.^[27,28]^ Yet, after administration, the administered material may deform or flatten under the stresses present in the subcutaneous space **(Figure 1a)**.^[29,30]^ This shape change has significant impact on diffusion length and corresponding release kinetics **(Figure 1c)**. The most common kinetic model used in drug release studies is the Ritger-Peppas model,^[31]^ which describes drug transport through both Fickian diffusion and anomalous transport mechanisms. Comparing Fickian release behaviors alone from an equal-volumed sphere and cylinder (aspect ratio eight) demonstrates that a spherical depot will result in a four-fold greater time to 60% release of the entrapped cargo **(Figure 1d)**. Additionally, a flattened cylinder has an increased surface area to volume ratio that would likely further increase the rate of erosion and corresponding drug release rate. Although simple modeling shows that shape heavily impacts drug release kinetics from polymeric drug delivery systems, little effort has focused on understanding and predicting spherical depot formation following hydrogel administration in the body. In this work, we seek to address this crucial gap in knowledge to elucidate how rheological properties of hydrogels are relevant to maintaining depot shape and persistence in the body.

**Figure 1.**
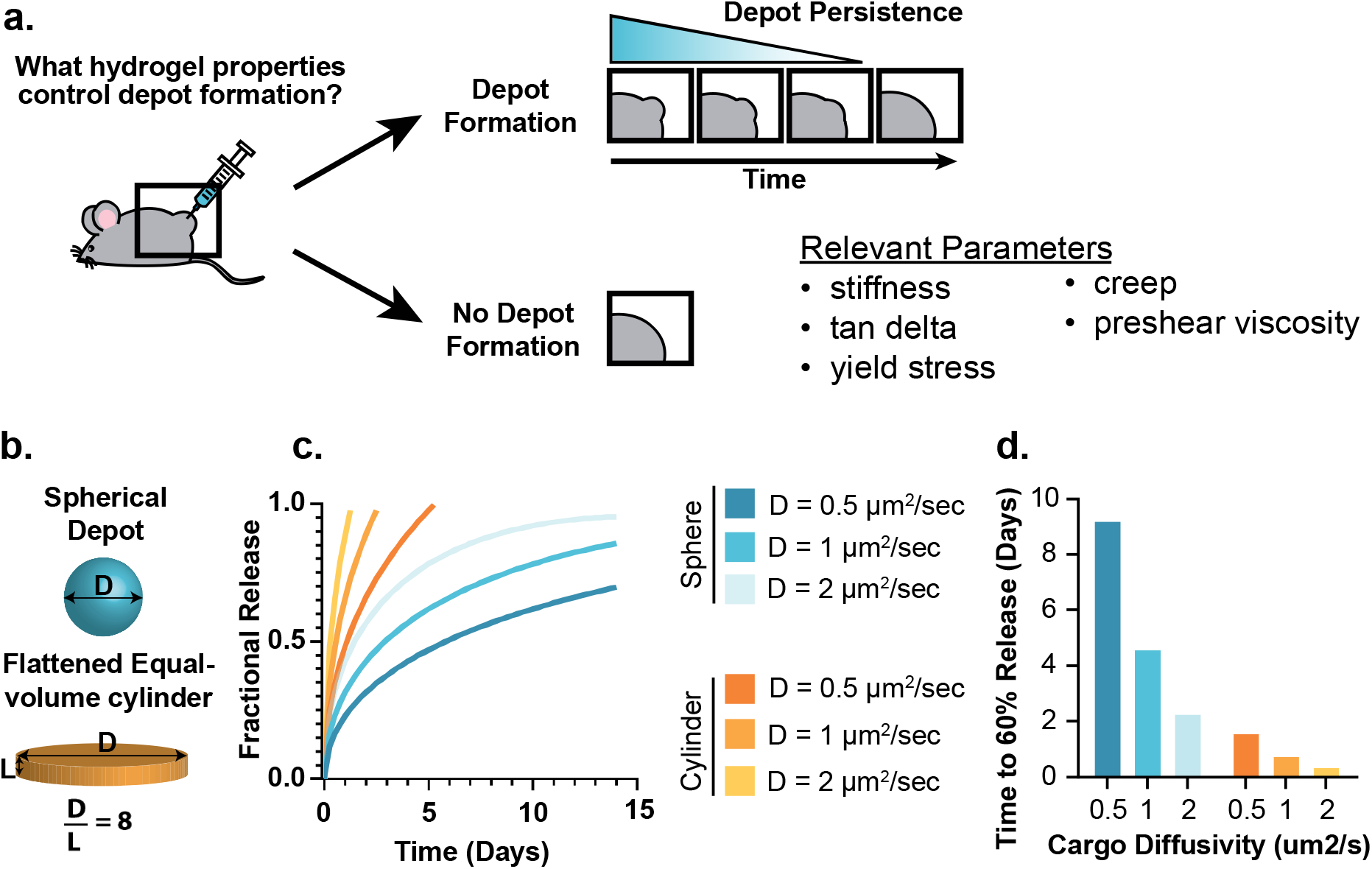
Schematic of injectable hydrogel depot formation and persistence. **a**. Illustration of depot persistence as compared to rapid flattening following subcutaneous administration. **b**. Illustration of spherical and cylindrical hydrogel depots. **c**. Cargo release profiles from spherical and cylindrical depots with varied diffusion coefficients as predicted by the Ritger-Peppas model. **d**. Impact of depot geometry and diffusion coefficient on time to 60% release of the entrapped cargo.

To probe the impact of gel rheological properties on depot formation and persistence, we utilize two physiochemically-distinct, physically crosslinked hydrogel systems, including calcium-alginate and polymer-nanoparticle (PNP) hydrogels **(Figure 2)**. First, PNP hydrogels are self-assembled from dynamic, multivalent, and entropically-driven non-covalent interactions between nanoparticles and high-molecular-weight biopolymers **(Figure 2a)**.^[ 32 ]^ We have previously reported PNP hydrogels formulated with poly(ethylene glycol)-b-poly(lactic acid) nanoparticles (PEG-PLA NPs) and dodecyl-modified hydroxypropylmethylcellulose polymers (HPMC-C_12_),^[33]^ whereby the HPMC-C_12_ polymers form a dynamic corona that surrounds and bridges between nanoparticles upon mixing of the two components.^[34]^ These materials are shear thinning and injectable and have been successfully utilized for various biomedical applications including prolonged delivery of therapeutic molecules and cells, postoperative adhesion barriers, stabilization of biopharmaceuticals, and prolonged delivery of wildland fire retardants for wildfire prevention.^[8,35,36,37,38]^ Second, calcium-alginate hydrogels are formed through the crosslinking of sodium alginate by calcium ions **(Figure 2d)**. The crosslinking of occurs through exchange of sodium ions from guluronic acid residues that make up segments of the polysaccharides with multivalent cations such as calcium or barium in aqueous media.^[39]^ On account of their simple crosslinking chemistry, high water content and resemblance to soft tissue, calcium-alginate hydrogels have been used in a multitude of therapeutic molecule and cell delivery, wound dressing, tissue engineering, and bioprinting applications.^[40,41,42,43,44]^

**Figure 2.**
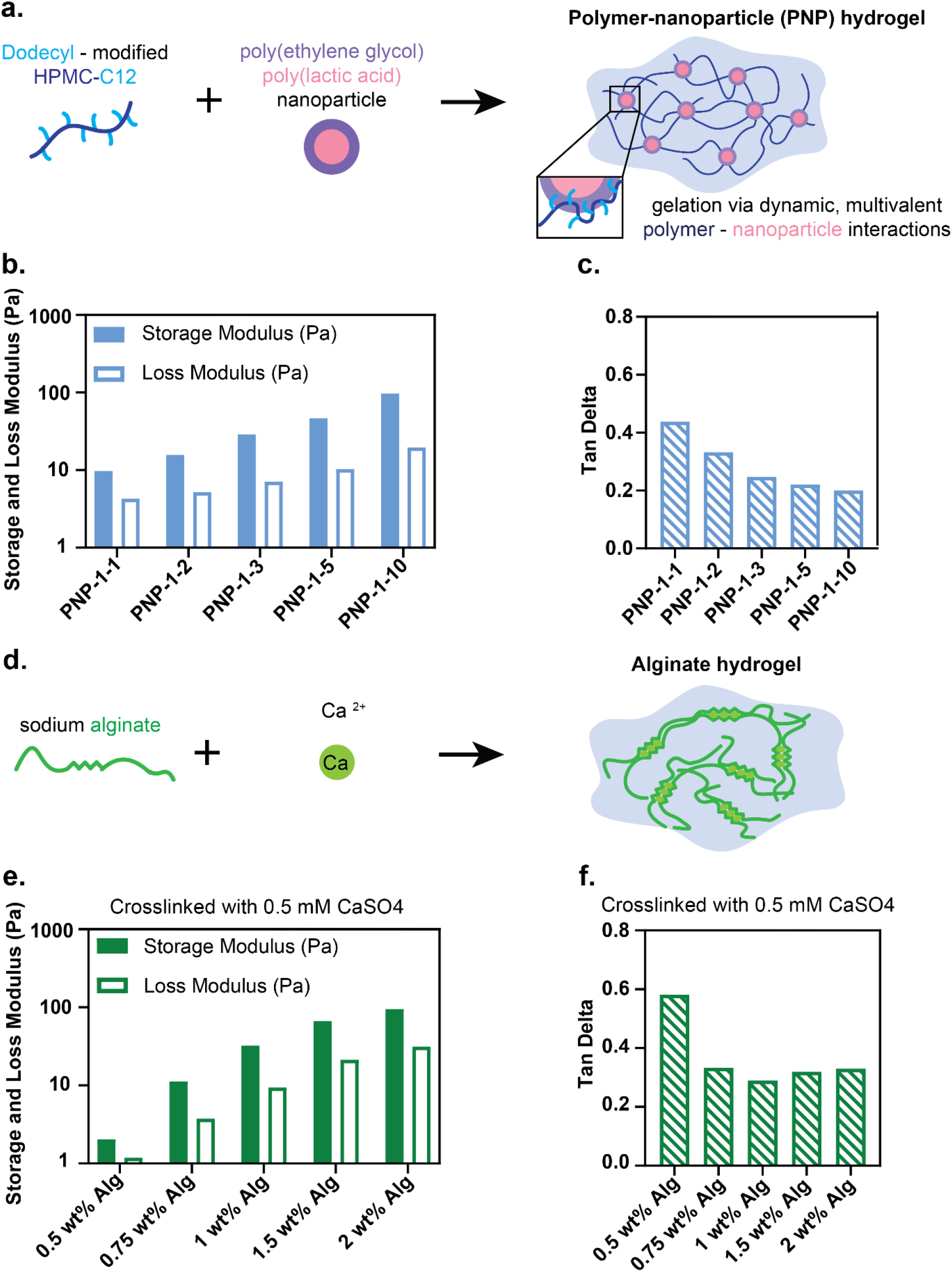
Two distinct non-covalently crosslinked hydrogels have tunable rheological properties. **a**. Formulation of polymer-nanoparticle hydrogel. **b**. Storage and loss moduli of PNP hydrogel formulations. **c**. Tan delta of PNP hydrogel formulations. **d**. Formulation of alginate hydrogel. **e**. Storage and loss moduli of alginate hydrogel formulations. **f**. Tan delta of alginate hydrogel formulations.

The rheological properties of both calcium-alginate and PNP hydrogels can be easily tuned. For example, either increasing the concentration of alginate at a constant calcium content or increasing the concentration of PEG-PLA NPs at a constant HPMC-C_12_ concentration^[45]^ yields hydrogels with more solid-like properties, higher yield stresses, higher preshear viscosities, and decreased creep **(Figure 2b, e)**. By thoroughly characterizing the mechanical properties of these materials and monitoring their behavior upon subcutaneous administration in mice, we aim to elucidate relationships between rheological behaviors and the ability to form and maintain depots following injection that can yield generalizable design criteria for injectable hydrogels.

## 2. Results and Discussion

### 2.1. Rheological Characterization of PNP and Alginate Hydrogels

To capture a range in gel rheological properties, we created five PNP hydrogels with a uniform HPMC-C_12_ polymer content of 1 wt% and varied PEG-PLA NP contents of 1, 2, 3, 5, and 10 wt%. PNP hydrogel formulations are referred to in the format P-NP, whereby P denotes the weight percent of HPMC-C_12_ and NP denotes the weight percent of the PEG-PLA NPs. The remaining mass of the formulation is phosphate-buffered saline. As the PEG-PLA NPs act as crosslinkers in the PNP hydrogel system, increasing NP content in these materials results in hydrogels with more solid-like properties as indicated by the increase in storage modulus and decrease in tan-delta **(Figure 2b, c)**. Furthermore, NP content impacted both yield stress and preshear viscosity. Stress-controlled yield stress measurements evaluate hydrogel viscosity while slowly increasing the stress the hydrogel is exposed to **(Figure 3a)**. At stresses below the hydrogel’s yield stress, the material does not flow and exhibited a high preshear viscosity. When the stress exceeds the hydrogel’s yield stress, the material starts to flow, and the viscosity was observed to drop significantly. As NP content of PNP hydrogels increases, the crosslink density increases and resulted in an increase in both the material’s yield stress and preshear viscosity prior to yielding **(Figure 3b)**.

**Figure 3.**
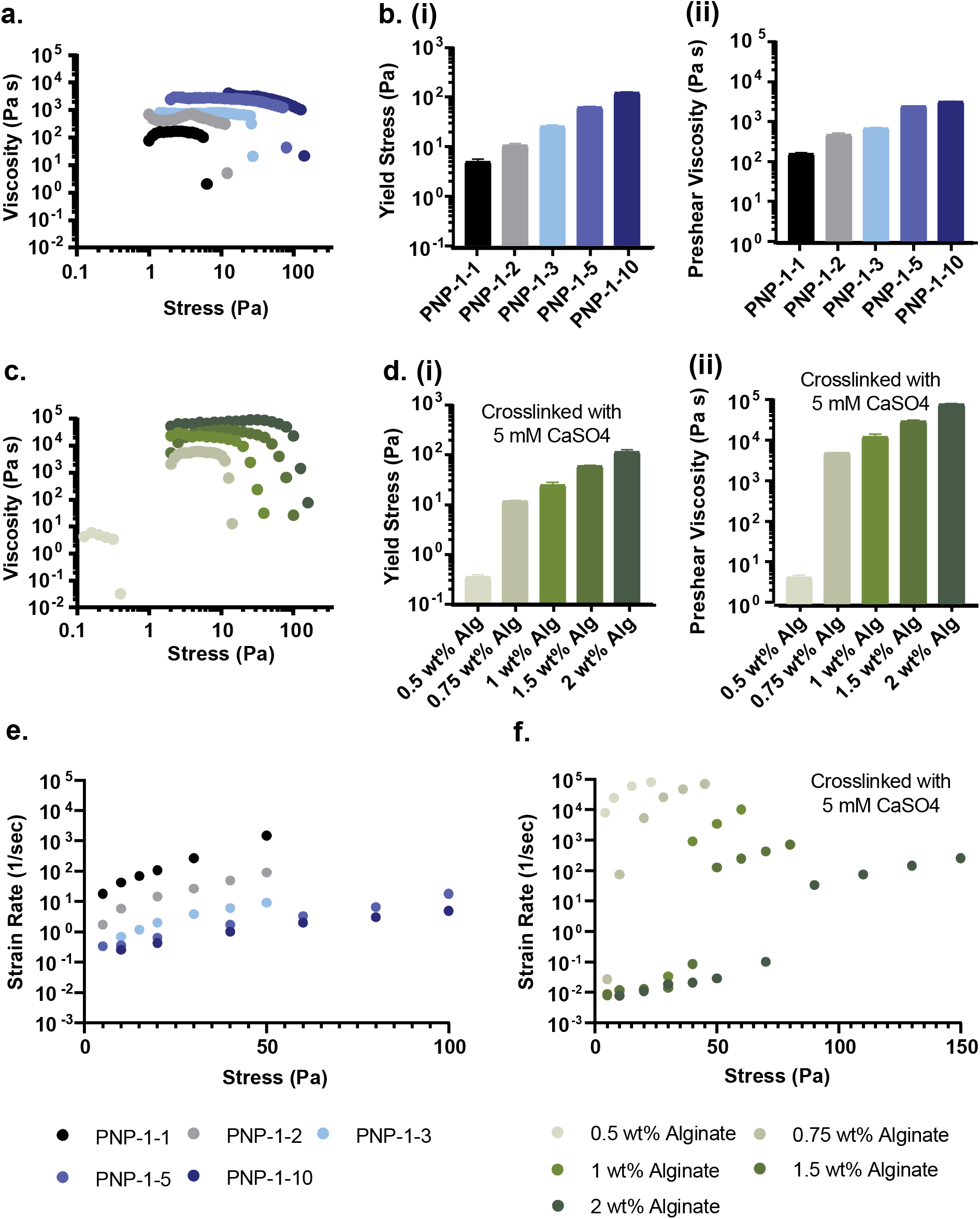
PNP and Alginate hydrogels exhibit tunable yield stress, preshear viscosity, and creep behavior. **a**. Stress-controlled yield stress measurement of PNP hydrogels. **b**. PNP hydrogel **(i)** yield stress and **(ii)** pre-shear viscosity. **c**. Stress-controlled yield stress measurement of calcium-alginate hydrogels. **d**. Calcium-alginate hydrogel **(i)** yield stress and **(ii)** pre-shear viscosity. **e**. Creep performance of PNP hydrogels. **f**. Creep performance of calcium-alginate hydrogels.

Calcium-alginate hydrogels exhibited similar rheological trends as alginate polymer content is increased. We created five distinct calcium-alginate hydrogels with a uniform calcium sulfate concentration of 0.5mM and varied alginate weight percent of 0.5, 0.75, 1, 1.5 and 2 wt% alginate. As alginate content increased, these hydrogels showed increased yield stress and increased preshear viscosity prior to yielding **(Figure 3d)**. It is interesting to note that while PNP hydrogels and alginate hydrogels exhibited nearly matched yield stresses, calcium-alginate hydrogels showed much higher preshear viscosities.

Creep is an additional rheological property that we hypothesized to be relevant to depot persistence time following subcutaneous administration. A creep test studies the deformation of a material following exposure to a small but constant stress. PNP hydrogel formulations exhibited lower strain rates at a given stress with increased NP content, and the observed strain rate within a given formulation increases log-linearly with stress **(Figure 3e)**. Calcium-alginate hydrogels exhibited distinct creep behavior, whereby strain rate below the yield stress was small, but exhibited a significant increase above the yield stress. Moreover, the strain rate observed above the yield stress is lower for stiffer materials comprising higher alginate content **(Figure 3f)**. The unique creep behavior of the PNP and alginate hydrogels likely arises from large differences in the rates of association and disassociation of the physical crosslinks in these systems. PNP hydrogels have highly dynamic crosslinks that break and reform rapidly even when the hydrogels are exposed to stresses exceeding its yield stress. Calcium-alginate hydrogels behave more similarly to covalently crosslinked materials, whereby the crosslinks remain intact below the yield stress when the material is not flowing, but break apart above the yield stress, resulting in a rapidly straining material unable to recover its network structure.

### 2.2. Depot Formation and Persistence Following Injection in the Subcutaneous Space

We hypothesized that hydrogel depot formation and persistence following subcutaneous administration can be predicted by hydrogel rheology. Specifically, we expected yield stress to be predictive of depot formation and creep behavior to be predictive of depot persistence. To probe this question, we subcutaneously injected fluorescently tagged calcium-alginate and PNP hydrogel formulations (100 μL) exhibiting variable mechanical behaviors across a range of rheological properties (stiffness, tan delta, yield stress, and creep). We then utilized brightfield photographic images collected with a standard camera and fluorescent images collected from an In-Vivo Imaging System (IVIS) to study depot formation and persistence at the site of injection.

#### 2.2.1. The Role of Hydrogel Yield Stress in Predicting Depot Formation

If the yield stress of the hydrogel is less than the stresses exerted by the tissue in the subcutaneous space, we would expect the hydrogel to flow and flatten. In contrast, if the yield stress of the hydrogel is greater than these stresses, we would expect the hydrogel to maintain its shape and form a robust, roughly spherical depot. We observed with both PNP (**Figure 4a**) and calcium-alginate (**Figure 5a**) hydrogels that there exists a minimum hydrogel yield stress needed to prevent immediate flattening after administration in the subcutaneous space. Hydrogels with yield stresses less than or equal to 12 Pa flatten shortly after administration while hydrogels with yield stresses greater than or equal to 25 Pa persist as stable depots for more than two weeks. From these observations, we estimate the stress of the subcutaneous space of a mouse to be between 12 and 25 Pa. Additionally, although the depots of low yield stress hydrogels appear to flatten shortly after administration, it is evident from fluorescent IVIS imaging of the injection site that the hydrogel components are still present at the site of injection two weeks later, though with poorly defined shape **(Figure 4b, 5b)**.

**Figure 4.**
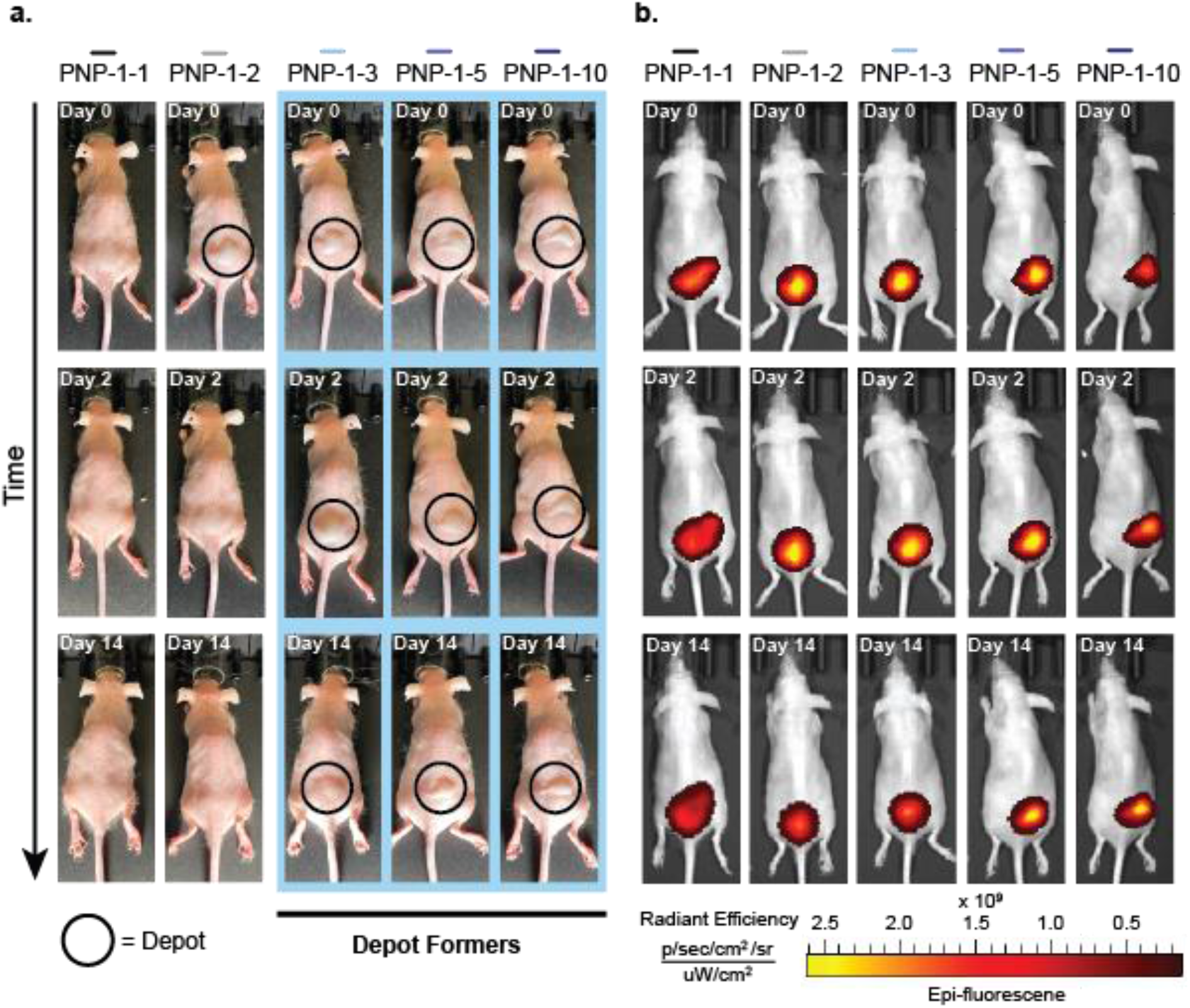
PNP gel rheological properties impact depot formation and persistence. **a**. Photographic images of PNP hydrogel depots at 0, 2, and 14 days after injection. **b**. IVIS images of PNP hydrogel depots at 0, 2, and 14 days after injection.

**Figure 5.**
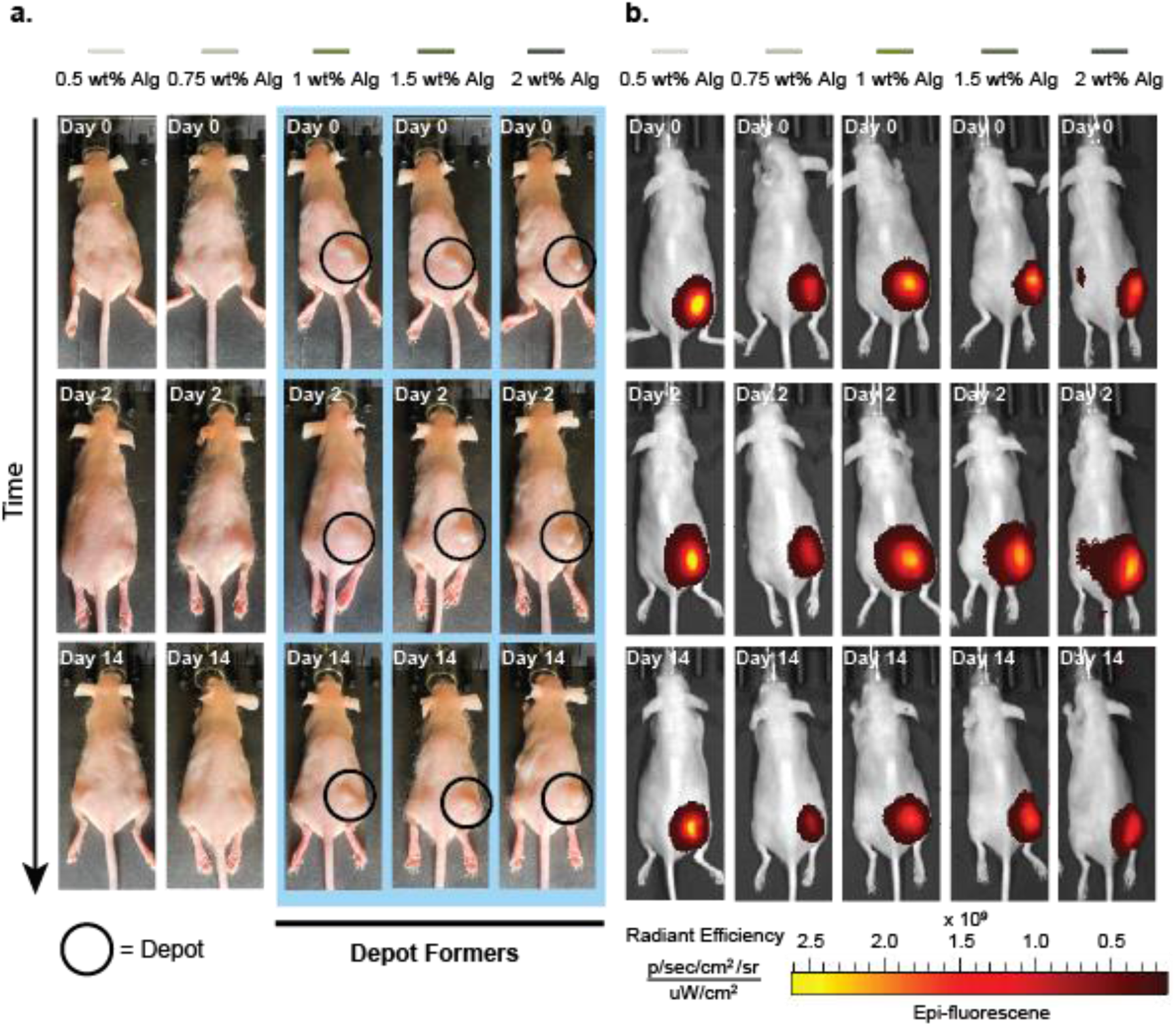
Alginate gel rheological properties impact depot formation and persistence. **a**. Photographic images of calcium-alginate hydrogel depots at 0, 2, and 14 days after injection. **b**. IVIS images of calcium-alginate hydrogel depots at 0, 2, and 14 days after injection.

#### 2.2.2. Quantification of Depot Persistence

For both PNP and calcium-alginate hydrogels, formulation dramatically impacts depot persistence time whereby stiffer hydrogel formulations persist for a longer time (**Figure 6a**). PNP-1-1, PNP-1-2, PNP-1-3, PNP-1-5, and PNP-1-10 formulations persisting for 0.4, 2.9, 17.4, 22.6, and 24.4 days, respectively. Depot persistence time of the depot-forming PNP hydrogel formulations (e.g., PNP-1-3, PNP-1-5, PNP-1-10) was found to be statistically greater than the depot persistence time of non-depot forming formulations (e.g., PNP-1-1, PNP-1-2) with p<0.0001.

**Figure 6.**
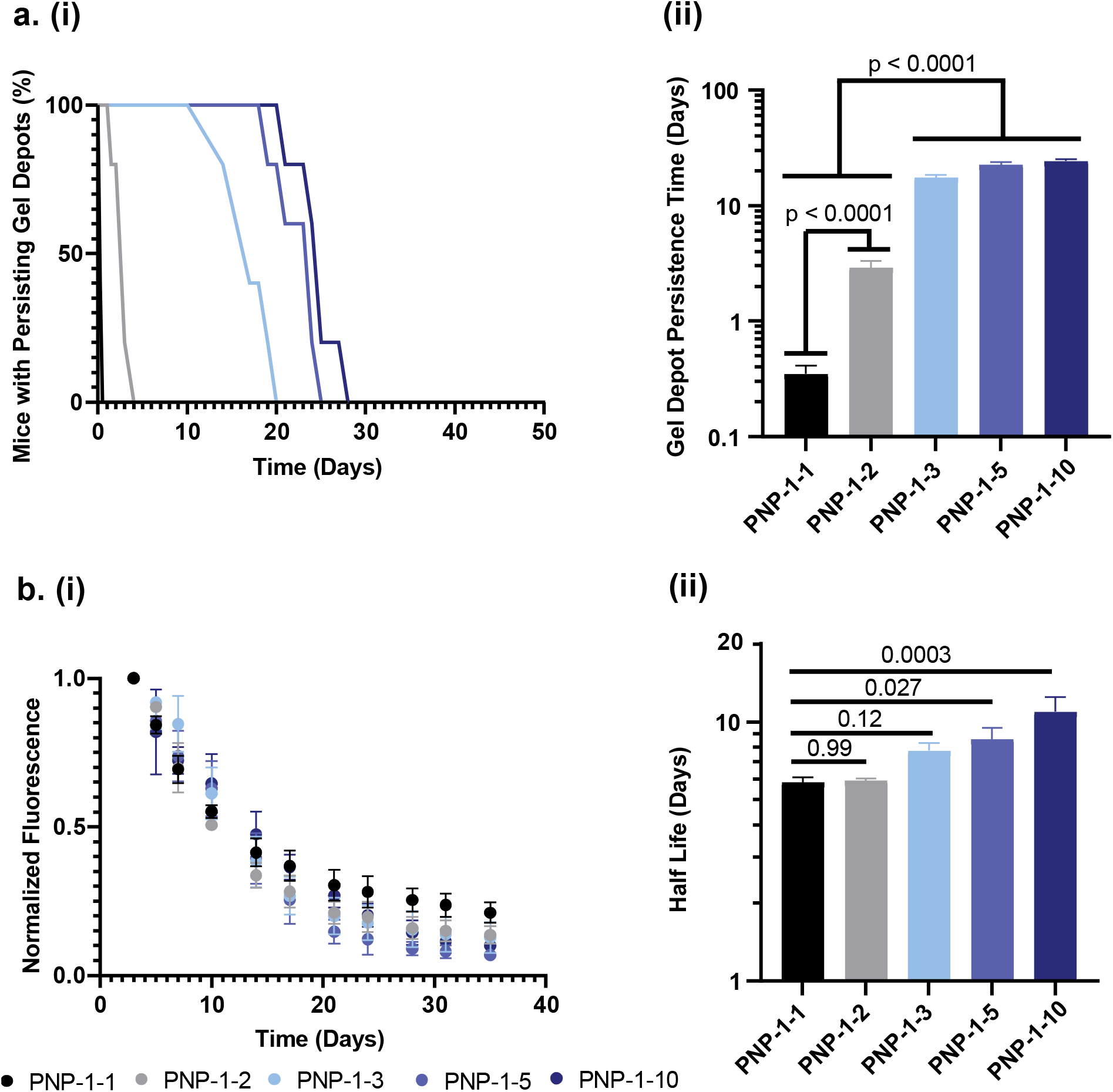
Increasing nanoparticle content in PNP hydrogels increases depot persistence time and half-life of gel retention in the subcutaneous space. **a. (i)** Fraction of mice with persisting gel depots over time from each PNP formulation and **(ii)** average depot persistence time for each PNP formulation. **b. (i)** Curves illustrating average fluorescent signal decay from PNP gels and **(ii)** half-lives calculated by fitting fluorescent decay curves to a single-phase exponential decay model.

IVIS imaging of fluorescently tagged PNP hydrogels provides further insight into gel retention in the subcutaneous space. Fluorescent signal over time from each gel injection was normalized and fit to a single-phase exponential decay curve to provide a half-life of gel retention (**Figure 6b**). Stiffer PNP hydrogel formulations showed increased half-lives of hydrogel retention with PNP-1-1, PNP-1-2, PNP-1-3, PNP-1-5, and PNP-1-10 formulations exhibiting average half-lives of 5.8, 5.9, 7.7, 8.5, and 10.9 days respectively. The increase in half-life of hydrogel retention for stiffer formulations that form more robust depots likely results from decreased rates of hydrogel erosion.

Similar trends in depot persistance are observed for calcium-alginate hydrogels, whereby stiffer calcium-alginate gel formulations persist for a longer time (**Figure 7a**). Formulations comprising 0.5 wt% alginate, 0.75 wt% alginate, 1 wt% alginate, 1.5 wt% alginate, and 2 wt% alginate persisted for 0.3, 1.3, 26, 29, and 37 days, respectively. As some calcium-alginate hydrogels failed to flatten completely due to a deleterious immune response, we report the median hydrogel depot persistence time rather than the mean (**Figure S3**).

**Figure 7.**
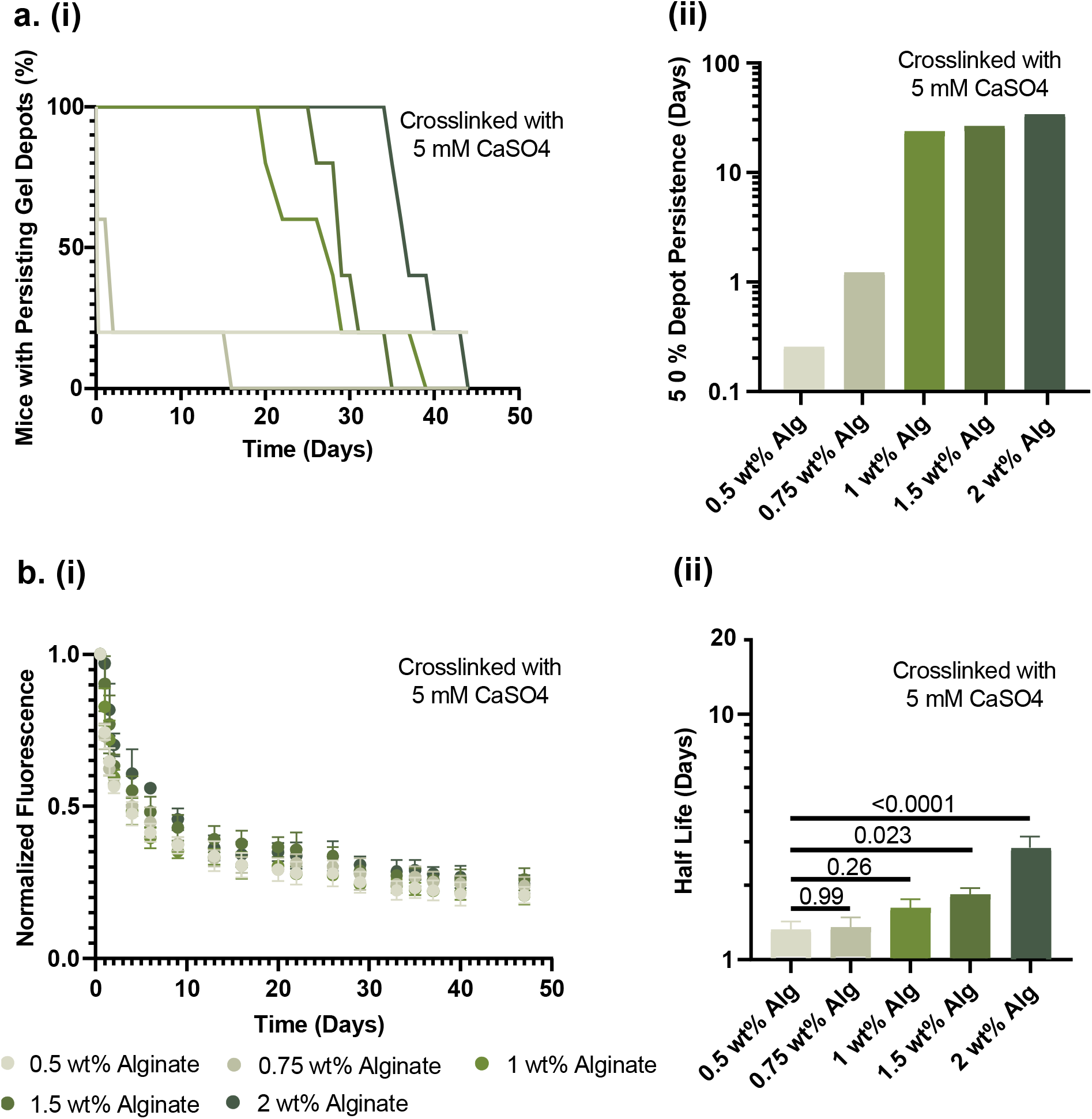
Increasing polymer content in alginate hydrogels increases depot persistence time and half-life of gel retention in the subcutaneous space. **a. (i)** Fraction of mice with persisting hydrogel depots over time from each calcium-alginate formulation and **(ii)** time when only 50% of calcium-alginate depots in each hydrogel group persist. **b. (i)** Curves illustrating average fluorescent signal decay from calcium-alginate hydrogels and **(ii)** half-lives calculated by fitting fluorescent decay curves to a single-phase exponential decay model.

Stiffer calcium-alginate hydrogel formulations again showed increased half-lives of depot retention with 0.5 wt% alginate, 0.75 wt% alginate, 1 wt% alginate, 1.5 wt% alginate, and 2 wt% alginate formulations exhibiting average half-lives of 1.3, 1.4, 1.7, 1.9, and 3.1 days, respectively (**Figure 7b**). The shorter half-life fit for calcium-alginate hydrogels as comparted to PNP hydrogels likely results from calcium-alginate gels having a steeper initial fluorescence decay followed by a more gradual fluorescence decay at later timepoints (**Figure 7b(i)**). At 3 weeks, PNP hydrogels exhibited an average normalized fluorescence of 0.16 as compared to calcium-alginate hydrogels, which exhibited an average normalized fluorescence of 0.27. While the half-life of fluorescence decay for calcium-alginate hydrogels may be influenced by an early loss of fluorescence, overall fluorescence retention is longer for calcium-alginate hydrogels as compared to PNP hydrogels, corroborating the observed increase in depot persistence time.

#### 2.2.3. Materials Properties Contributing to Extended Depot Persistence

To identify rheological properties that are predictive of depot persistence time, we plotted hydrogel storage modulus, tan delta, yield stress, and preshear viscosity against the depot persistence time for each hydrogel formulation **(Figure 8)**. Although no single property fully predicts depot persistence time, it was apparent that persistence time of both hydrogel types increased with increasing storage modulus, yield stress, and preshear viscosity. Tan delta, a measure of relative elasticity of the hydrogels, does not appear to be universally predictive of depot persistence time. Indeed, PNP hydrogels showed a decrease in depot persistance time with increasing tan delta while calcium-alginate hydrogels showed an increase in depot persistence time with increasing tan delta.

**Figure 8.**
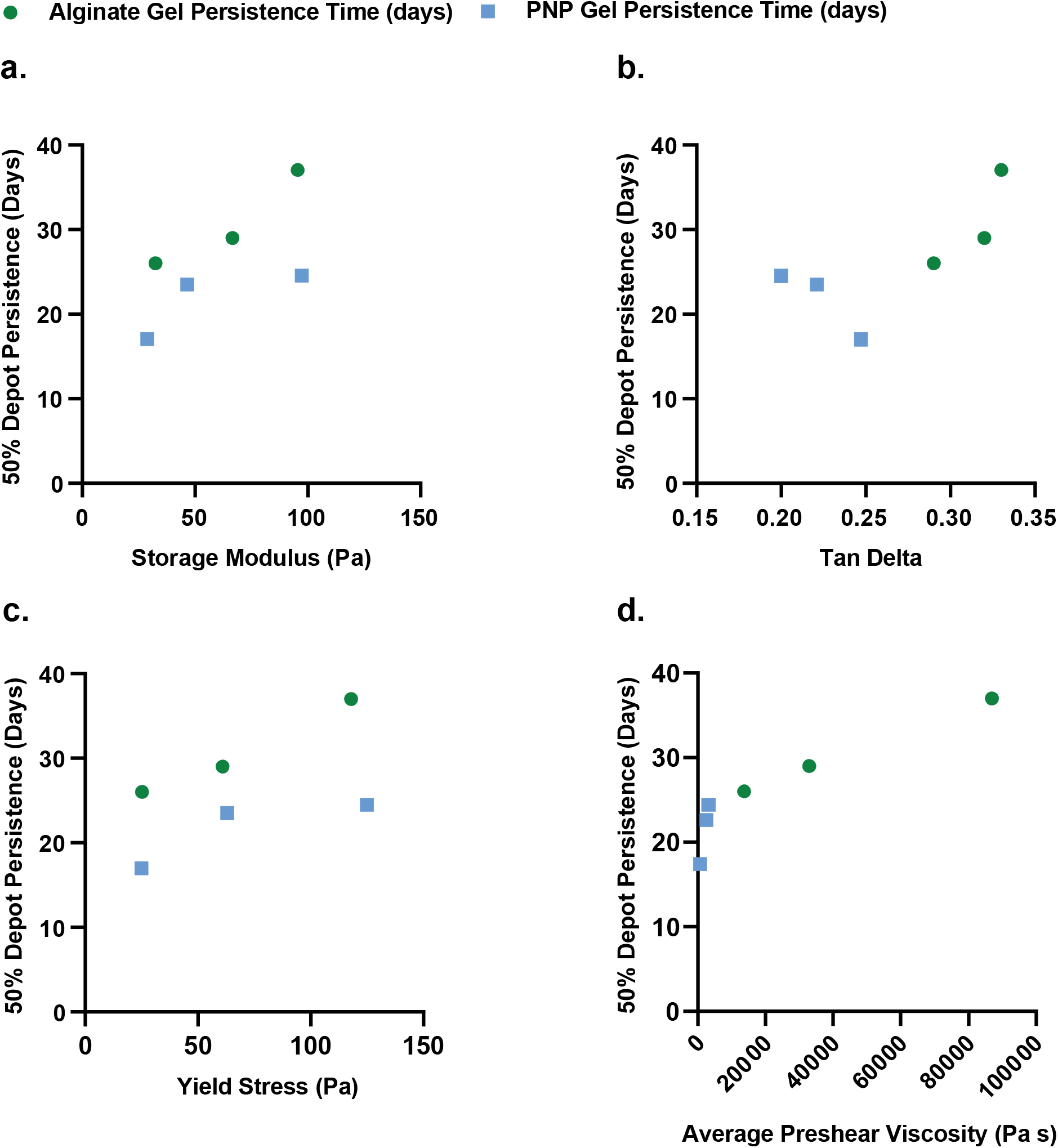
Storage modulus, yield stress, and viscosity are predictive of depot persistence time. Impact of **a**. storage modulus, **b**. tan delta, **c**. yield stress, and **d**. preshear viscosity on time to when only 50% of depots in each gel group persist.

While storage modulus, yield stress, and preshear viscosity are somewhat predictive of depot persistence time, they fail to explain the significantly higher depot persistence times of yield stress matched calcium-alginate hydrogels as compared to PNP hydrogels. One important way these hydrogel classes vary in their mechanical behavior is in their creep performance. We showed that the strain rates the hydrogels experience at a stress relevant to the subcutaneous space (e.g. 12-25 Pa), is highly predictive of relative depot persistence time. Indeed, the relative strain rates the hydrogels experience at a stress of 12-25 Pa is indicative of depot persistence time for both depot-forming and non-depot-forming, yield-stress-matched PNP and calcium-alginate hydrogels. For non-depot-forming hydrogels with a yield stress of 12 Pa, calcium-alginate gels exhibit a significantly higher strain rate than PNP hydrogels at a stress relevant to the subcutaneous space, and these calcium-alginate materials exhibit a commensurate decrease in depot persistence time than their PNP hydrogel counterparts **(Figure 9a)**. In contrast, for depot-forming hydrogels with yield stresses of 25, 60, and 120 Pa, calcium-alginate gels exhibit a significantly lower strain rate at a stress relevant to the subcutaneous space than PNP hydrogels, and these calcium-alginate materials exhibit a commensurate increase in depot persistence time compared to their PNP hydrogel counterparts **(Figure 9b-d)**.

**Figure 9.**
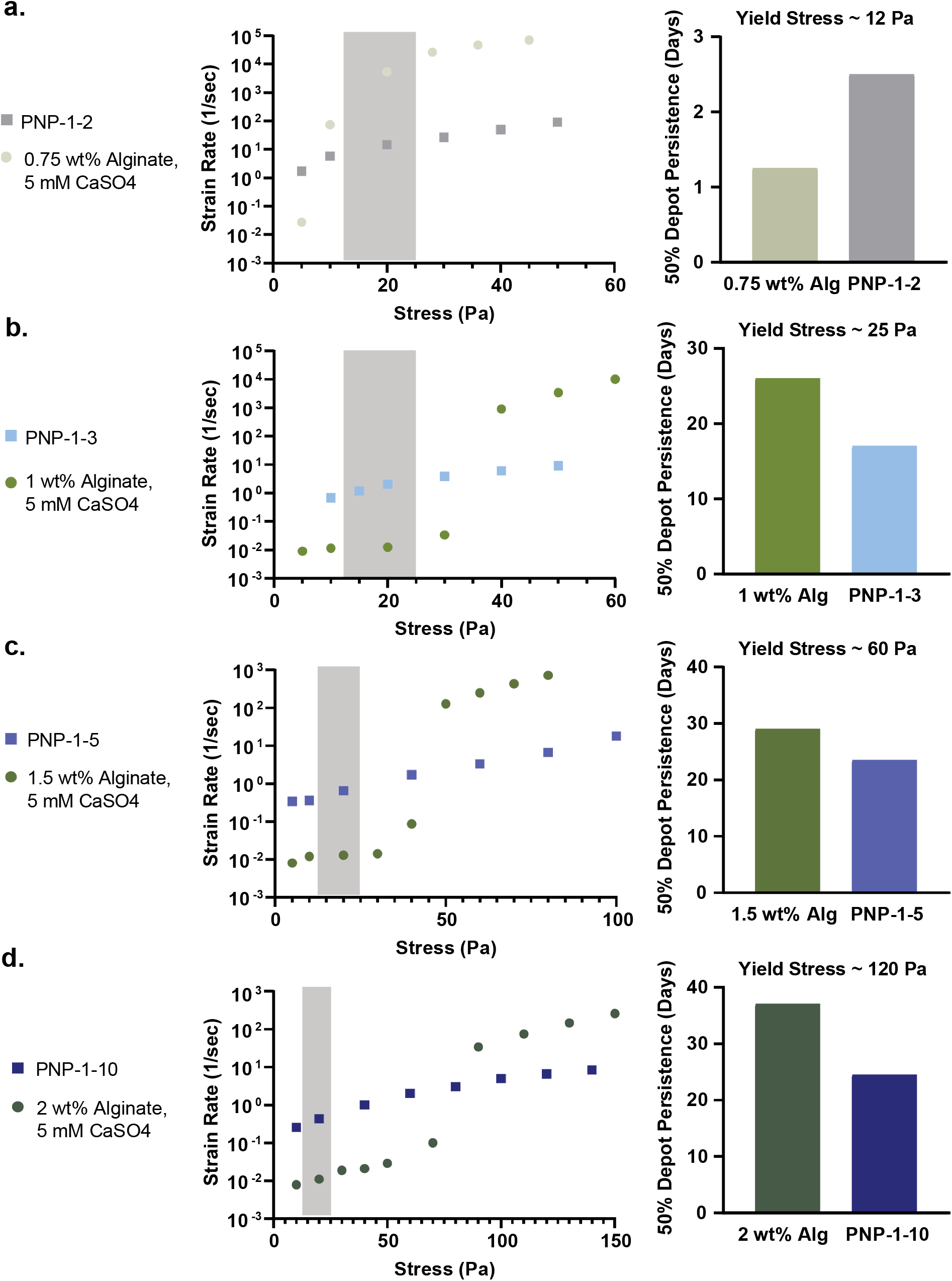
Relative strain rate at a stress relevant to that experienced in the subcutaneous space is predictive of depot persistence time. Graphs illustrating relationship between strain rate and applied stress as well as depot persistence time for yield-stress matched PNP and alginate hydrogels with yield stresses of **a**. 12 Pa, **b**. 25 Pa, **c**. 60 Pa, and **d**. 120 Pa.

This observation is further quantified by plotting the relationship between strain rate at a stress relevant to the subcutaneous space and depot persistence time. We observe a log-linear relationship between the strain rate observed from creep tests at 15 Pa and depot persistence time **(Figure 10a)**. Depot-forming and non-depot-forming hydrogel formulations collapse onto separate log-linear curves, but each log-linear curve includes both physicochemically-distinct hydrogel classes. These observations suggest that creep performace of physical hydrogel materials may be useful in predicting depot persistence time *in-vivo*, which is highly relevant to numerous biomedical applications. As viscosity is a materials property frequently extracted from creep test data, we also plotted the relationship between average pre-shear viscosity obtained from stress-controlled yield stress measurements and depot persistence time **(Figure 10b)**. As preshear viscosity is only relevant to gels forming a robust depot, only depot-forming hydrogel formulations were included in this analysis. Again, a log-linear relationship between pre-shear viscosity and depot persistence time was observed that captured both calcium-alginate and PNP hydrogel materials, suggesting that pre-shear viscosity may have predictive power in estimating depot persistence times. Future work will be directed at confirming these trends with additional hydrogel chemistries to both assign physical meaning to the observed log-linear relationship and generate generalizable design criteria for physical hydrogel materials.

**Figure 10.**
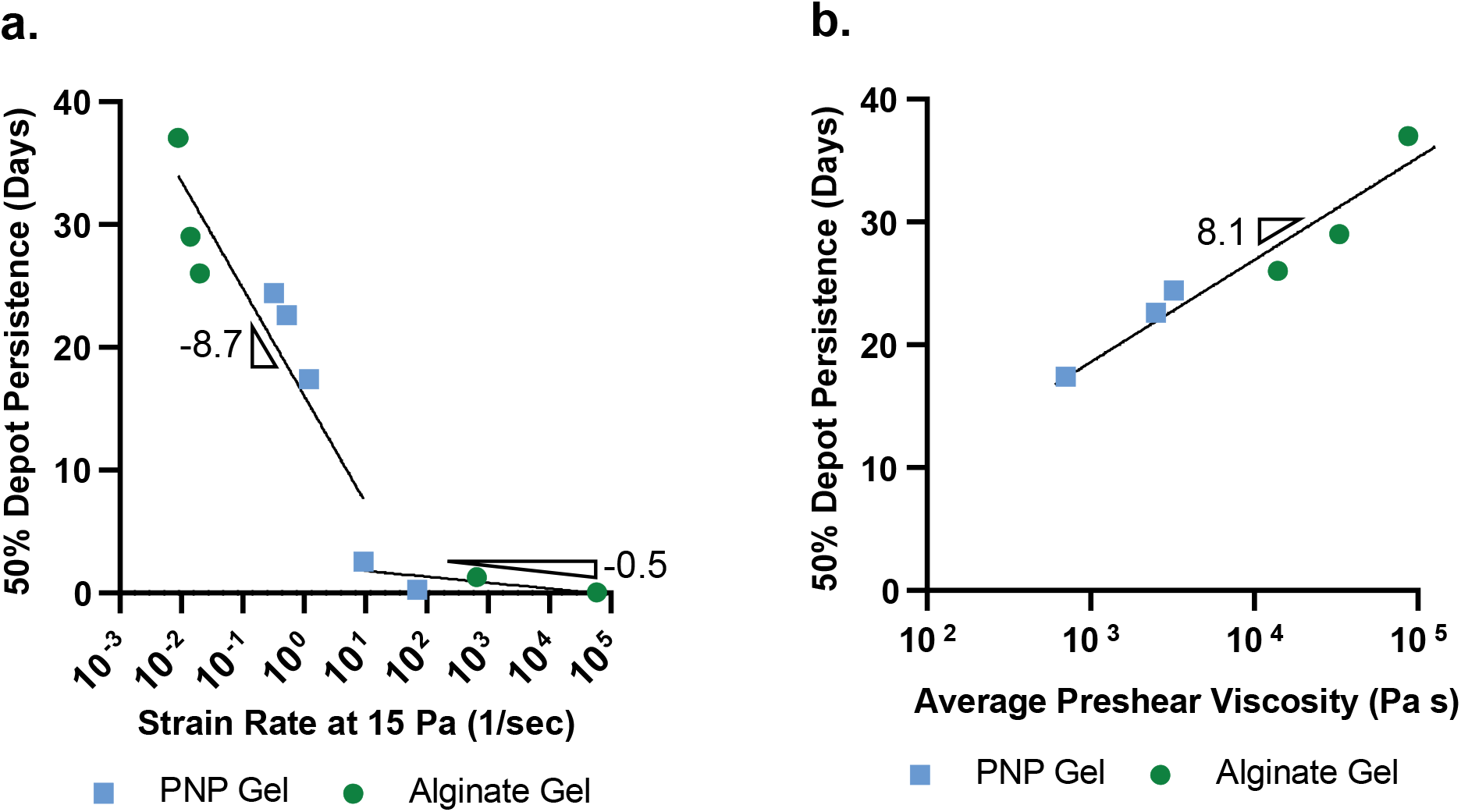
Strain rate and preshear viscosity are predictive of depot persistence time. **a**. Log-linear relationship between strain rate at 15 Pa and time when only 50% of depots in each gel group persist. **b**. Log-linear relationship between preshear viscosity and time when only 50% of depots in each gel group persist.

## 3. Conclusion

Understanding how hydrogel properties affect depot formation following administration in the body is crucial to the development of controlled release technologies. In this work, we evaluated how hydrogel rheology can be used to predict depot formation and persistence following subcutaneous administration in mice. We utilized both calcium-alginate hydrogels and PNP hydrogels, which are two physicochemically-distinct, physically crosslinked hydrogel systems. These hydrogel systems are formulated to exhibit variable mechanical behaviors across a range of important rheological properties (stiffness, tan delta, yield stress, and creep), and the added advantage of calcium-alginate and PNP hydrogels with matched yield stresses allowed for the decoupling of the impacts of yield stress and creep behaviors. By relating measured rheological properties to depot persistence time when injected into the subcutaneous space of mice, we identify that yield stress is predictive of initial depot formation while creep is predictive of depot persistence. Hydrogels with yield stresses greater than 25 Pa form robust depots, and depot persistence time is log-linearly related to the strain rate values obtained from creep tests. These findings provide predictive insights into design considerations for hydrogel technologies capable of extended controlled release of therapeutic cargo.

## 4. Experimental Section

### 4.1 Materials

Sodium alginate was purchased from Bioworld (CAS:9005-38-3). Racemic lactide (3,6-dimethyl-1,4-dioxane-2,5-dione, 99%, Sigma-Aldrich) was recrystallized twice from dry ethyl acetate to remove impurities, with sodium sulfate included as a desiccant prior to the first recrystallization. Dichloromethane (DCM) was dried immediately before use by cryogenic distillation. HPMC (USP grade), N,N-Diisopropylethylamine (Hunig’s base), hexanes, diethylether, N-methyl-2-pyrrolidone (NMP), 1-dodecylisocynate, diazobicylcoundecene (DBU), 2-(N-morpholino) ethanesulfonic acid (MES), 1-Ethyl-3-(3 dimethylaminopropyl) carbodiimide (EDC), and N-hydroxysulfosuccinimide (NHS) sulfo ester were purchased from Sigma Aldrich and used as received. Monomethoxy-PEG and Monomethoxy-PEG-azide (5 kDa) were purchased from Sigma Aldrich and purified by azeotropic distillation with toluene prior to use. AF647-DBCO was purchased from Thermo Fisher and used as received. Sulfo-Cyanine7 amine was purchased from Lumiprobe and used as received. All other materials were purchased from Sigma-Aldrich and used as received.

### 4.2. Methods

#### Poly(ethylene)-block-poly(lactic acid) synthesis

Poly(ethylene glycol)-block-poly(lactic acid) (PEG-b-PLA) block copolymers were synthesized by ring-opening polymerization of lactide onto a PEG macromer, as described previously.^[33]^ Dry PEG methyl ether (*M*_n_ = 5000 g/mol; 0.5 g, 0.1 mmol) and 1,8-diazabicyclo[5.4.0]undec-7-ene (DBU, 98%; 15 μL, 0.1 mmol) were dissolved together in dry DCM (1.5 mL); twice recrystallized racemic lactide (2 g, 13.9 mmol) was dissolved separately in dry DCM (9 mL). The PEG/DBU solution was quickly added to the lactide solution under nitrogen atmosphere and stirred rapidly for 8 min then quenched with dilute acetic acid in acetone (100 μL). The polymer was recovered by precipitating twice from a 50:50 diethyl ether: hexanes solution and dried under vacuum. Target molecular weight of the block copolymers by DMF SEC: 22.5–27.5 kDa (5 kDa PEG, 17.5–22.5 kDa PLA with Đ < 1.2)

#### PEG-b-PLA nanoparticle synthesis

To form core–shell NPs, PEG–PLA (50 mg) was dissolved in a 75:25 acetonitrile:DMSO solution (1 mL) and added dropwise to ultrapure water (10 mL) at a high stir rate (600 rpm). The NP solution was centrifuged over a filter (Amicon Ultra-15, threshold molecular weight 10 kDa) and resuspended in PBS as a 20 wt % stock solution (total volume 250 μL per batch). The resulting NPs were characterized by dynamic light scattering (DLS, DynaPro II plate reader, Wyatt Technology) (hydrodynamic diameter = 30-35 nm; PDI < 0.2).

#### Fluorescent PEG-b-PLA nanoparticle synthesis

AF647 NPs were prepared using a combination of PEG–PLA (25 mg) and unconjugated azide-PEG–PLA (25 mg). A 1 ml solution of combined PEG–PLA and azide-PEG–PLA in DMSO (50 mg/ml) was added dropwise to 10 ml of water at room temperature under a high stir rate (600 rpm). NPs were purified by ultracentrifugation over a filter (molecular weight cut-off of 10 kDa; Millipore Amicon Ultra-15) followed by resuspension in water to a final concentration of 200 mg/ml. The nanoparticles were then functionalized by mixing adize-functional NPs (250 μL, 20 wt%) with AF647-DBCO (25 μl, 1 mg/ml) and waiting 12 hours.

#### Dodecyl-modified hydroxypropylmethylcellulose (HPMC-C12) synthesis

Dodecyl-modified (hydroxypropyl)methyl cellulose (HPMC) was synthesized as previously described.^[33]^ Briefly, HPMC (hypromellose, USP grade, 1 g) was dissolved in 1-methyl-2-pyrrolodinone (NMP, 40 mL) at 80 °C. Dodecyl isocyanate (99%, 125 μL, 0.52 mmol) dissolved in NMP (5 mL) was added dropwise to the heated reaction vessel followed by 10 drops of *N,N*-diisopropyethylamine (≥99%, Hünig’s base) catalyst. Reaction solution was allowed to cool to room temperature and stirred overnight. HPMC-C_12_ was recovered by precipitation in acetone and purified by dialysis (MWCO 3500 Da) for 4 days. Lyophilized HPMC-C_12_ was redissolved in phosphate buffered saline (PBS) as a 6 wt % stock solution. Representative ^1^H NMR analysis has been previously published. ^[34]^

#### Polymer-nanoparticle hydrogel synthesis

Supramolecular PNP hydrogels were formulated by combining the HPMC-C_12_ stock solution and PEG-PLA NP stock solution diluted to the desired concentration via elbow mixing as previously described.^[36]^ PNP hydrogel formulations are designated by HPMC-C_12_ wt % - NP wt %. For example, a PNP-1-10 hydrogel comprises 1 wt % HPMC-C_12_ and 10 wt % NPs (i.e., 11 wt % total solids) of the total formulation mass, with the remainder comprised of PBS. HPMC-C_12_ stock solution (167 mg) was loaded into a 3 mL luer lock syringe. NP stock solution (500 μL) was combined with PBS (333 μL) and loaded into a 3 mL luer lock syringe. The HPMC-C_12_-filled syringe was connected to a female luer × female luer elbow fitting, and the solution was pushed through the elbow to eliminate air in the connection before attaching the NP-filled syringe to the other side. The solutions were mixed through the elbow at a relatively fast rate for >100 cycles (1 cycle = complete transfer of material from one syringe to the other and then back into original syringe) until a homogeneous hydrogel was formed. All other PNP hydrogel formulations (PNP-1-1, PNP-1-2, PNP-1-3, PNP-1-5) were prepared identically at the appropriate concentrations.

#### Fluorescent polymer-nanoparticle hydrogel synthesis

PNP hydrogels were made according to procedures above. Each 1000 μL of gel contained 50 μl of fluorescent AF647-conjugated PEG-PLA NPs and the remainder standard PEG-PLA NPs allowing for all gel formations to have matched fluorescence. For example, 1000 μL of PNP-1-10 contained 50 μl of fluorescent AF647-conjugated PEG-PLA NPs and 450 μL of standard PEG-PLA NPs.

#### Alginate hydrogel synthesis

Calcium-alginate hydrogels were formulated by crosslinking sodium alginate with CaSO_4_. Sodium alginate (Bioworld CAS:9005-38-3) was dissolved in PBS at a 4 wt% solution. CaSO_4_ was added to PBS at a concentration of 250 mM to form a slurry. The 250 mM CaSO_4_ slurry and the alginate stock solution were combined with PBS to form calcium-alginate hydrogels. To form a hydrogel with 5 mM CaSO_4_ and 1 wt% alginate, the CaSO4 slurry (250 mM) was stirred rapidly and slurry solution (20 μL) was pipetted into PBS (730 μL) and loaded into a 3 mL luer lock syringe. Alginate stock solution (250 mg) was loaded into another 3 mL luer lock syringe. The CaSO_4_ filled syringe was connected to a female luer × female luer elbow fitting, and the solution was pushed through the elbow to eliminate air in the connection before attaching the alginate filled syringe to the other side. The solutions were mixed through the elbow at a relatively fast rate for >100 cycles (1 cycle = complete transfer of material from one syringe to the other and then back into original syringe) until a homogeneous hydrogel was formed. All other calcium-alginate hydrogel formulations (0.5 wt%, 0.75 wt%, 1.5 wt %, 2 wt% alginate) were prepared identically at the appropriate concentrations.

#### Florescent tagging of alginate

Sodium alginate (Bioworld CAS:9005-38-3) (150 mg) was dissolved in 0.1 M 2-(N-morpholino) ethanesulfonic acid (MES) buffer (15 mL) at a pH of 6 and stirred for 1.5 hours to ensure complete dissolution. SulfoCy7 amine (2.5 mg), 1-Ethyl-3-(3-dimethylaminopropyl) carbodiimide (EDC) (72.5 mg), and N-hydroxysulfosuccinimide (NHS) sulfo ester (41 mg) were then added. The mixture was stirred for 20 hours and transitioned from a green to dark blue color. It was then dialyzed against MilliQ water for 5 days (MWCO 3.5 kDa) and lyophilized.

#### Fluorescent alginate hydrogel synthesis

Alginate hydrogels were made according to procedures above. Each 1000 μL of gel contained

0.5 wt% fluorescently tagged alginate and the remainder standard alginate allowing for all gel formations to have matched fluorescence. For example, 1000 μL of a hydrogel comprising 5 mM CaSO_4_ and 1 wt% alginate contained 125 mg of fluorescently tagged alginate and 125 mg of standard alginate.

#### Rheological Characterization

Rheological testing was performed using a 20 mm diameter serrated parallel plate at a 500 μm gap on a stress-controlled TA Instruments DHR-2 rheometer. All experiments were performed at 25 °C. Frequency sweeps were performed at a strain of 1% within the linear viscoelastic regime. Flow sweeps were performed from high to low shear rates with steady state sensing. Stress controlled yield stress measurements (stress sweeps) were performed from low to high stress with steady state sensing and 10 points per decade. Creep experiments measured strain rate at fixed stress. An initial stress of 0.5 Pa was applied to the material for 20 seconds. The desired stress (5 – 120 Pa) was then applied, and corresponding strain percent was measured for 2000 seconds. Strain rate at a given stress was obtained by fitting a slope for the linear region of the strain vs time curve in GraphPad Prism.

#### Animal Studies

All animal studies were performed in accordance with National Institutes of Health guidelines and with the approval of the Stanford Administrative Panel on Laboratory Animal Care (APLAC-32109).

#### In-vivo imaging of PNP gel depots

SKH1E mice were each administered 100 μL of PNP hydrogel via transcutaneous injection. Each of the five PNP formulations had a sample size of five mice, and PNP hydrogel formulations were each cage blocked. Mice were imaged using an In-Vivo Imaging System (IVIS Lago) over a series of timepoints spanning 35 days. When imaged, mice were anesthetized with isoflurane gas and imaged with an exposure time of 0.25 seconds, excitation wavelength of 600nm, and emission wavelength of 670 nm (binning: medium, F/stop: 1.2). Total radiant efficiency ([photons/s] / [μW/cm^2^]) was quantified using an equal-sized region of interest surrounding the gel depot. As early time points (time < 3 days) showed fluorescence in the region of interest to increase instead of decrease, fluorescent intensity at each timepoint was normalized to fluorescent intensity on day 3. Normalized fluorescence intensity values for each mouse (n=5) between day 3 and 35 were fit to a single exponential decay models and half-lives were acquired and averaged using GraphPad Prism.

#### In-vivo imaging of alginate gel depots

SKH1E mice were each administered 100 μL of alginate hydrogel via transcutaneous injection. Each of the five alginate formulations had a sample size of five mice, and alginate hydrogel formulations were each cage blocked. Mice were imaged using the in-vivo Imaging System (IVIS Lago) over a series of timepoints spanning 47 days. When imaged, mice were anesthetized with isoflurane gas and imaged with an exposure time of 2 seconds, excitation wavelength of 720 nm, and emission wavelength of 790 nm (binning: medium, F/stop: 1.2). Total radiant efficiency ([photons/s] / [μW/cm^2^]) was quantified using an equal-sized region of interest surrounding the gel depot. Fluorescent intensity at each timepoint was normalized to fluorescent intensity on day 0. Normalized fluorescence intensity values for each mouse (n=5) between day 0 and 30 were fit to a single exponential decay models and half-lives were acquired and averaged using GraphPad Prism.

#### Statistical Analysis

Animals were cage blocked, and Mead’s Resource Equation was used to identify a sample size above which additional subjects will have little impact on power. All post hoc tests were conducted with the Tukey HSD test in JMP and values presented are means and standard errors. Results were accepted as significant if p < 0.05.

## Data Availability Statement

The data that support the findings of this study are available on request from the corresponding author. The data are not publicly available due to privacy or ethical restrictions.

## Supporting Information

Supporting Information is available from the Wiley Online Library or from the author.

## Acknowledgements and Author Contributions

C.K.J. and E.A.A. conceived of the idea. C.K.J. performed experiments. Other authors aided with experiments. This research was financially supported by the Center for Human Systems Immunology with the Bill & Melinda Gates Foundation (OPP1113682), the Bill & Melinda Gates Foundation (OPP1211043; INV-027411), and the American Cancer Society (RSG-18-133-01). C.KJ. and J.Y. are thankful for National Science Foundation Graduate Research Fellowships. A.K.G. is appreciative of a National Science Foundation Graduate Research Fellowship and the Gabilan Fellowship of the Stanford Graduate Fellowship in Science and Engineering. J.B. is grateful for support from a Marie-Curie fellowship from the European Union under the program H2020, Grant 101030481.

