## Supplemental Figures for "Yield-Stress and Creep Control Depot Formation and Persistence of Injectable Hydrogels Following Subcutaneous Administration"

.

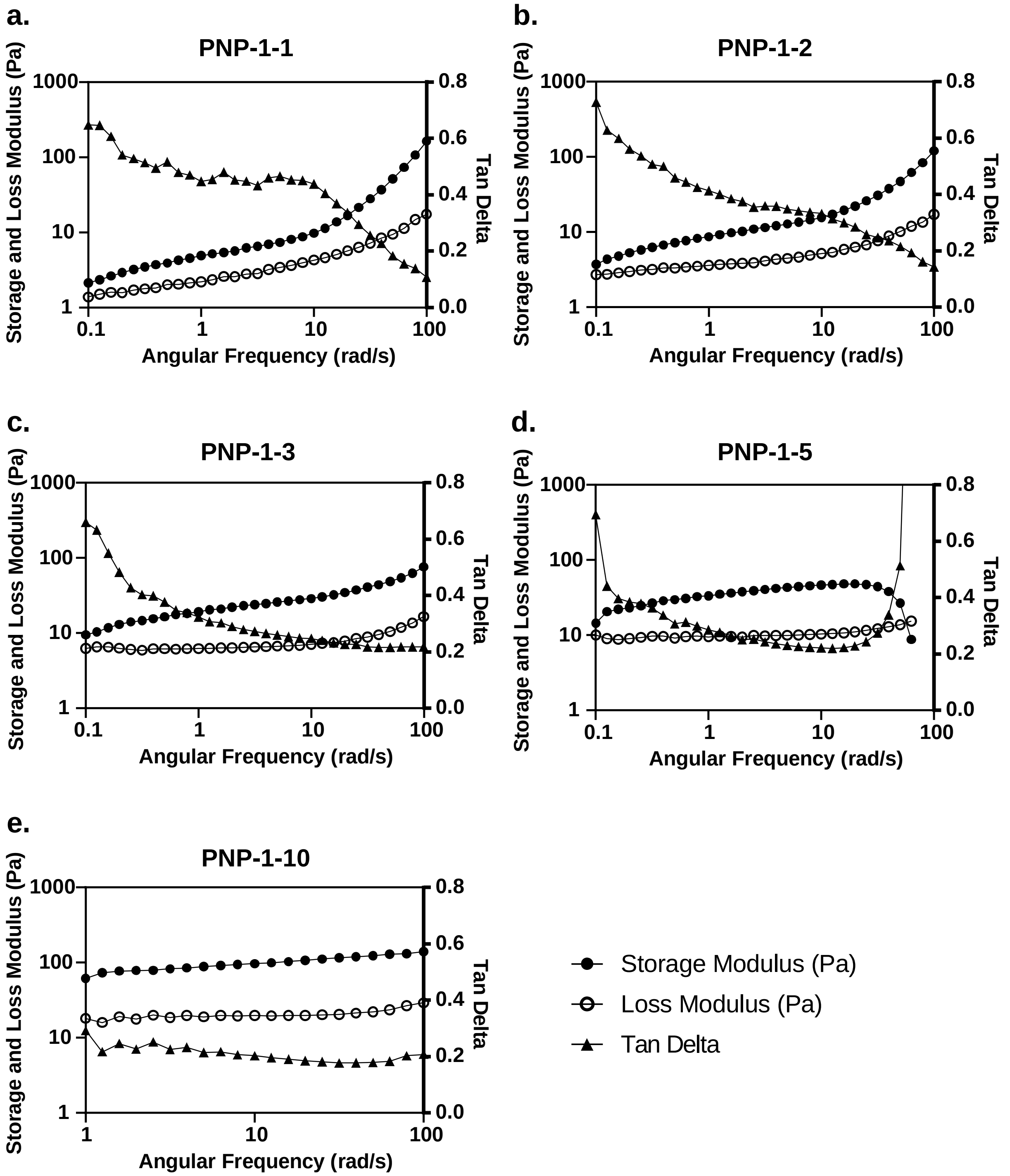

**Figure S1. Frequency sweeps characterize storage modulus, loss modulus, and tan delta of PNP hydrogels.** Frequency sweeps of **a.** PNP-1-1, **b.** PNP-1-2, **c.** PNP-1-3, **d.** PNP-1-5, and **e.** PNP-1-10 formulations performed at a strain of 1% within the linear viscoelastic regime.

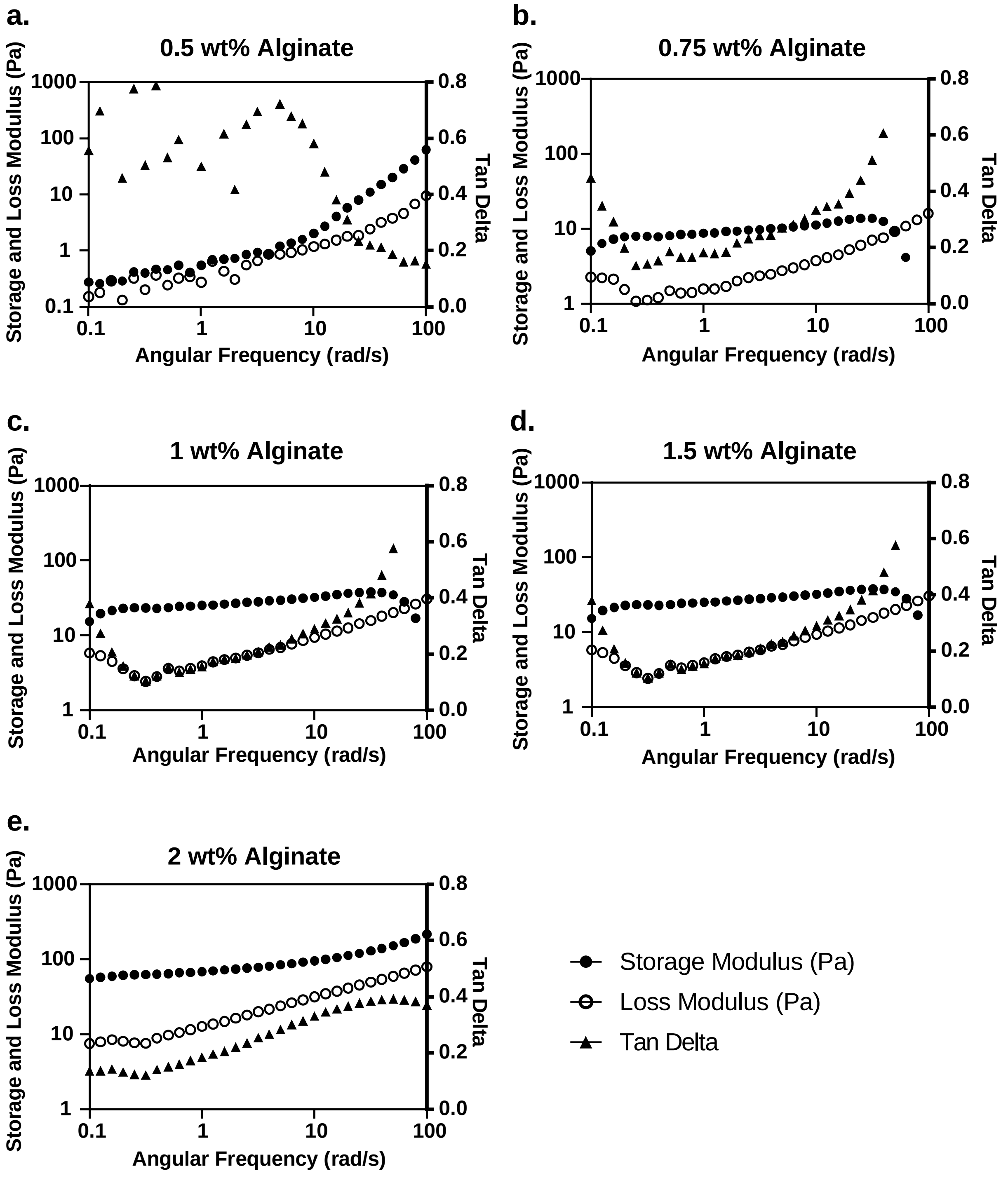

**Figure S2. Frequency sweeps characterize storage modulus, loss modulus, and tan delta of alginate hydrogels.** Frequency sweeps of **a.** 0.5 wt% alginate, **b.** 0.75 wt% alginate, **c.** 1 wt% alginate, **d.** 1.5 wt% alginate, and **e.** 2 wt% alginate formulations all crosslinked with 5mM $CaSO_4$. Frequency sweeps are performed at a strain of 1% within the linear viscoelastic regime.

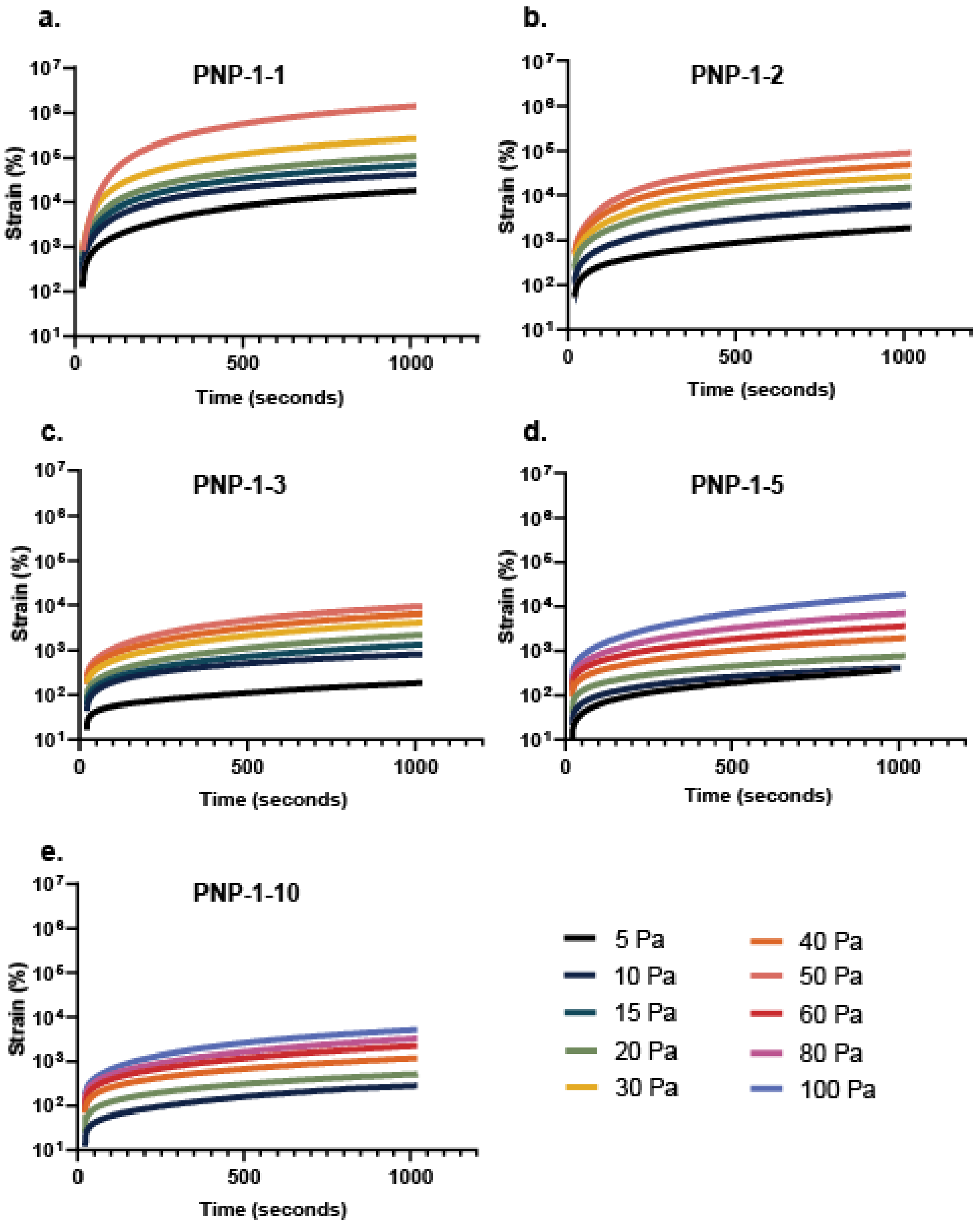

**Figure S3. Creep performance of PNP hydrogels.** Strain rates over time of **a.** PNP-1-1, **b.** PNP-1-2, **c.** PNP-1-3, **d.** PNP-1-5, and **e.** PNP-1-10 hydrogel formulations after a constant stress is applied.

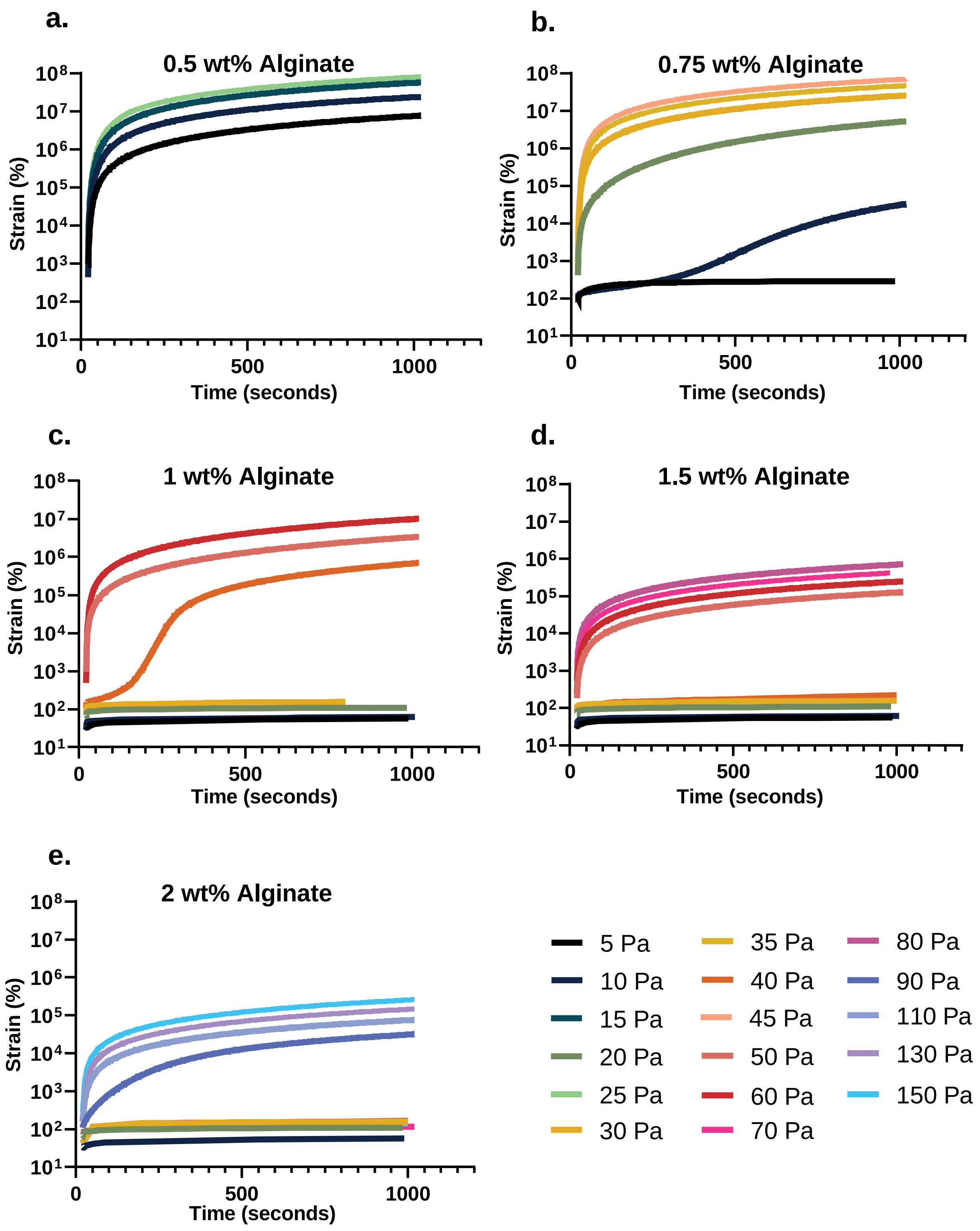

**Figure S4. Creep performance of alginate hydrogels.** Strain rates over time of **a.** 0.5 wt% alginate, **b.** 0.75 wt% alginate, **c.** 1 wt% alginate, **d.** 1.5 wt% alginate, and **e.** 2 wt% alginate hydrogel formulations after a constant stress is applied. All alginate hydrogel formations are crosslinked with 5mM $CaSO_4$.

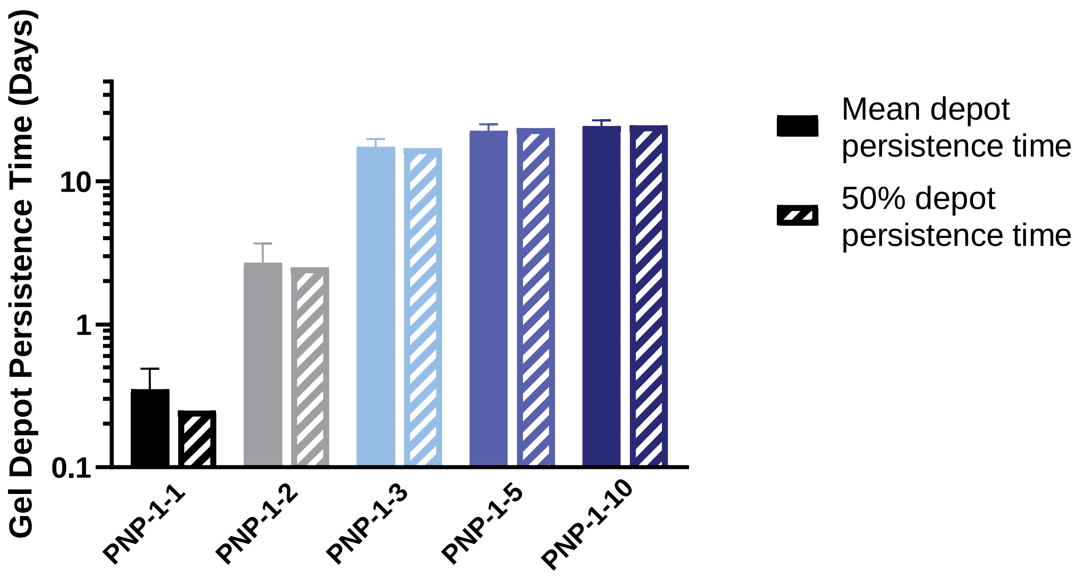

**Figure S5. Comparison of mean and 50% depot persistence time.** Measuring mean depot persistence time and time to 50% depot persistence results in comparable numerical values.
